# The multiple population genetic and demographic routes to islands of genomic divergence

**DOI:** 10.1101/673483

**Authors:** Claudio S. Quilodrán, Kristen Ruegg, Ashley T. Sendell-Price, Eric Anderson, Tim Coulson, Sonya Clegg

## Abstract

1. The way that organisms diverge into reproductively isolated species is a major question in biology. The recent accumulation of genomic data provides promising opportunities to understand the genomic landscape of divergence, which describes the distribution of differences across genomes. Genomic areas of unusually high differentiation have been called genomic islands of divergence. Their formation has been attributed to a variety of mechanisms, but a prominent hypothesis is that they result from divergent selection over a small portion of the genome, with surrounding areas homogenised by gene flow. Such islands have often been interpreted as being associated with divergence with gene flow. However other mechanisms related to genetic architecture and population history can also contribute to the formation of genomic islands of divergence.

2. We currently lack a quantitative framework to examine the dynamics of genomic landscapes under the complex and nuanced conditions that are found in natural systems. Here, we develop an individual-based simulation to explore the dynamics of diverging genomes under various scenarios of gene flow, selection and genotype-phenotype maps.

3. Our modelling results are consistent with empirical observations demonstrating the formation of genomic islands under genetic isolation. Importantly, we have quantified the range of conditions that produce genomic islands. We demonstrate that the initial level of genetic diversity, drift, time since divergence, linkage disequilibrium, strength of selection and gene flow are all important factors that can influence the formation of genomic islands. Because the accumulation of genomic differentiation over time tends to erode the signal of genomic islands, genomic islands are more likely to be observed in recently divergent taxa, although not all recently diverged taxa will necessarily exhibit islands of genomic divergence. Gene flow primarily slows the swamping of islands of divergence with time.

4. By using this framework, further studies may explore the relative influence of particular suites of events that contribute to the emergence of genomic islands under sympatric, parapatric and allopatric conditions. This approach represents a novel tool to explore quantitative expectations of the speciation process, and should prove useful in elucidating past and projecting future genomic evolution of any taxa.

## Introduction

A major aim of evolutionary biology is to understand mechanisms associated with the divergence of organisms between populations and the emergence of new species. This motivated Charles Darwin and Alfred Wallace 160 years ago when they advanced the Theory of Natural Selection (Darwin & Wallace 1858). Since then, the increasing accumulation of genetic, genomic and computational tools has allowed a better understanding of the genetic basis of the speciation process, resulting in the rise of a new era of evolutionary research (Hughes 2009; Chanderbali *et al*. 2016). Patterns of divergence at the level of the genome have been characterised for an increasing number of taxa, but the extent to which observed patterns are informative about evolutionary processes is actively debated (e.g. Ellegren *et al*. 2012; Renaut *et al*. 2013; Ruegg *et al*. 2014; Burri *et al*. 2015).

The genomic landscape of divergence describes the distribution of differences across the genomes of diverging organisms. The genome of a diverging taxon does not change uniformly, with some regions changing at higher rates than others (Seehausen *et al*. 2014; Ravinet *et al*. 2017). If a single process uniquely generates a particular divergence pattern, then identification of that pattern can confidently be interpreted as representing a particular evolutionary history. In contrast, if multiple processes can generate the same patterns of genomic divergence, then identification of the pattern will not point to a specific process, though the suite of candidate processes may be narrowed. In these cases, additional information beyond patterns of genomic divergence, such as the ecological and evolutionary context of a given divergence, will be required to understand patterns of evolutionary divergence.

Genomic islands of divergence – highly differentiated regions of the genome that are surrounded by regions of low differentiation – are a particularly intriguing pattern of genomic divergence (Turner, Hahn & Nuzhdin 2005; Harr 2006; Nosil, Funk & Ortiz – Barrientos 2009). Initially, their formation was attributed to the action of divergent natural selection on particular loci, creating elevated regions containing the selected loci and other physically linked loci, surrounded by regions homogenised by gene flow (Wu 2001; Wu & Ting 2004; Nosil, Funk & Ortiz – Barrientos 2009). Several authors consequently interpreted presence of genomic islands of divergence as a signal of divergence with gene flow (e.g. Feder *et al*. 2013), concluding that the speciation process in sympatric or parapatric conditions may be more common than previously thought (e.g. Nosil 2008; Fraïsse *et al*. 2014; Soria-Carrasco *et al*. 2014). However, empirical studies have also proposed that genomic islands of divergence can arise in the absence of gene flow due to a variety of causes, such as the architecture of the diverging genomes (e.g. variation in recombination rate) and the action of genetic drift, background selection, and adaptation to local environmental conditions (Noor & Bennett 2009; Cruickshank & Hahn 2014; Campagna *et al*. 2015). Furthermore, regions of low divergence may occur because of incomplete lineage sorting rather than homogenisation by gene flow, with peaks of genetic divergence being an artefact of a loss of nucleotide diversity after divergent selection (Cruickshank & Hahn 2014).

There have been some previous attempts to model the dynamic of the architecture of genomic landscapes that can be applied to the formation of genomic islands of divergence. However, these models either simulate single bi-allelic selected loci (Charlesworth, Nordborg & Charlesworth 1997; Sedghifar, Brandvain & Ralph 2016) or consider a small number of simulated loci (Feder & Nosil 2009; Feder & Nosil 2010; Feder *et al*. 2012). They also provide a static view of the divergence process by summarizing selection as a single parameter. While informative, such models represent specific stages when populations have already achieved a given level of differentiation. Flaxman, Feder and Nosil (2013) used an individual-based model to project this dynamic forward in time, but their model was constrained to a uniform distribution of loci with constant recombination rates. A quantitative and more flexible framework than previous attempts is thus required to evaluate the dynamics of genomic landscapes, and increase the utility of accumulated genomic datasets (Feder *et al*. 2013; Seehausen *et al*. 2014). We develop a quantitative individual-based modelling approach to simulate the dynamic of a genomic landscape of divergence. The model simulates any number of loci and allelic polymorphisms and can be theoretically motivated or parameterised using data. A major difference between our approach and previous simulations is the treatment of the fitness function, which is compatible with many structured ecological and evolutionary models. Our approach is highly flexible and can be constructed for any genotype-phenotype map and any configuration of recombination rates between neighbouring loci, can be constructed for deterministic and stochastic environments, and incorporates any desired system of mating. The simulation method presented here provides a flexible framework to examine the dynamics of diverging genomic landscapes under various scenarios of gene flow and selection on single genes or networks of multiple interacting genes. Our approach represents a novel tool to evaluate quantitative expectations in genomic landscapes. It is useful to elucidate the influence of a range of demographic and evolutionary scenarios, including divergence with or without gene flow, the divergence timeframe, and the architecture of target genomes.

## Methods

### General description of the model

The purpose of our model is to provide insight into how a range of genetic and demographic processes can generate genomic signatures and patterns of genomic divergence between populations. Our primary motivation was to explore factors associated with the emergence of genomic islands of divergence, but our approach can be applied to many questions about genetic architectures, genomic landscapes, and the evolution of divergent organisms.

The model is individual-based and consists of two populations that may or may not be linked by gene-flow. Our model is composed of three hierarchical levels: genotypes, phenotypic traits, and demographic rates. The dynamics of the populations, the distributions of genotypes at each locus and the phenotypic traits, are all emergent properties of the model. The model tracks the multivariate distribution of multi-locus genotypes and phenotypes. We simulate individuals that are characterised by sex and genetic identity (Fig. 1a). The genotype and the environment determine the phenotypic trait values of an individual via a genotype-phenotype map. The phenotypic trait values influence an individual’s expected demography (i.e. survival, mate choice, and reproductive success). For example, assuming a per generation time step, the potential number of offspring produced by each individual depends on its phenotype *ω* = *f* (*z*), which in turn depends on the individual genotype and on the environment *z* = *g*(*G, E*) (Fig. 1b). *G* is a numeric value determined by an individual’s genotype, representing the genetic value of the genotype. In the case of an additive genetic map, the genetic value of a genotype will be a breeding value. *E* represents the effect of the environment on phenotypic expression, and this allows us to capture the effects of plasticity on phenotypic expression. The environmental effect is important when simulating real-life eco-evolutionary dynamics because it almost always interacts with the genotype to determine the expression of a phenotypic trait (Bradshaw 1965; Kokko *et al*. 2017). The realized demography is obtained by sampling from a distribution whose expected value is the expected demography. Once mating pairs are formed, the genotype of the young is determined by merging haploid gametes produced by each parent. Genetic variation of the offspring is determined by recombination and mutation.

**Fig. 1.**
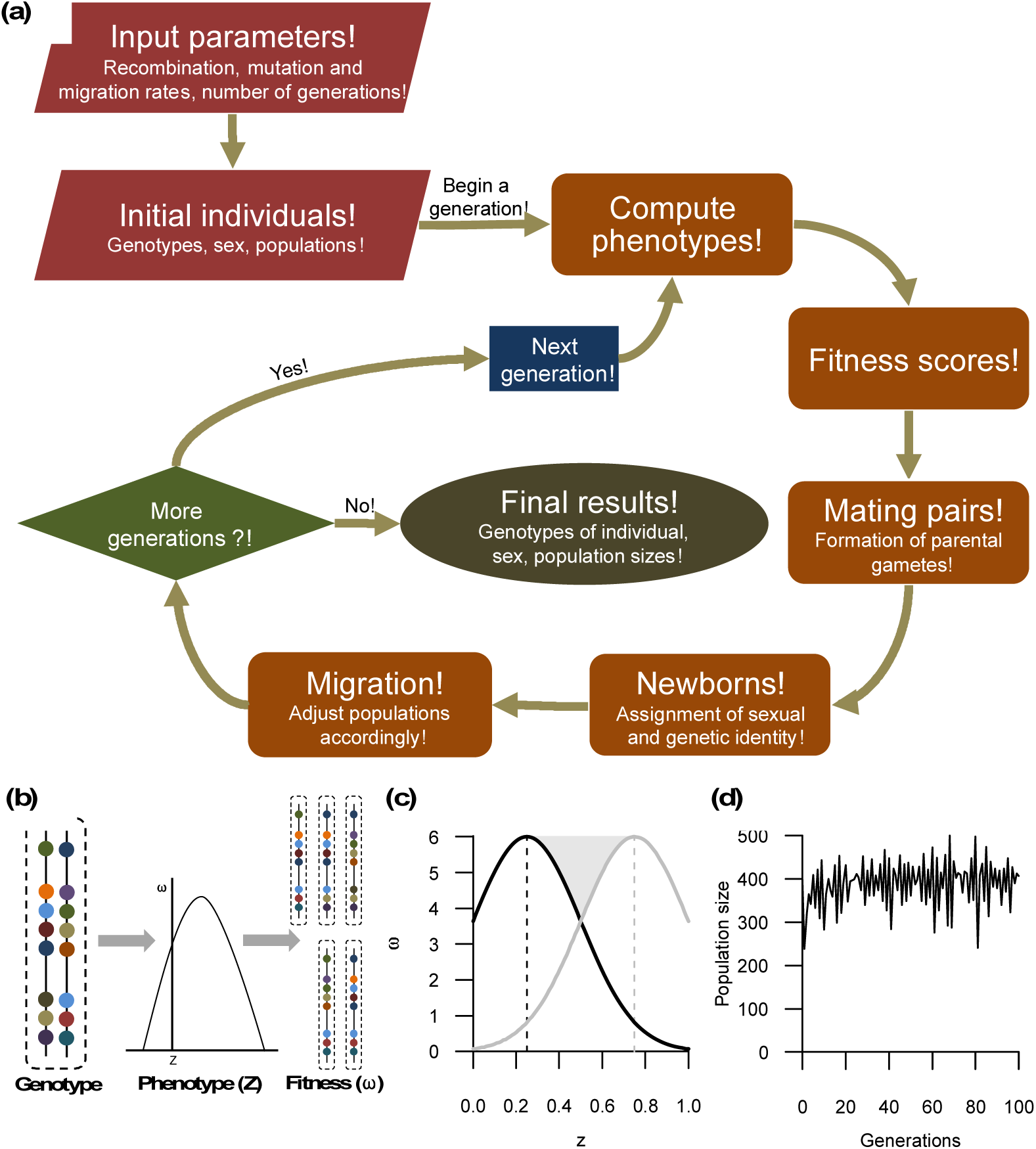
General description of the simulation framework. A) Main steps of the general modelling approach. The red polygons represent the starting conditions. The orange squares are the different computing steps on each generation. The green polygon is a condition variable stating either the running of a next generation (blue square) or the end of simulations (olive green circle). B) Relationship between genotypes, phenotypes, and fitness. The genetic variation is represented in different colours. The space between points represents unequal centiMorgan distances. C) Two different fitness functions with different phenotypic optima. D) Example of population size across time.

We describe how the model is implemented in the next section. Our starting point is the distribution of individuals classified by genotype, sex and population (Fig. 1a). First, we generate the phenotypic trait of each individual given its genotype (and potentially the environment). Second, we calculate individual fitness given an individual’s phenotype, the population it is in, and potentially the environment. Mating pairs are formed based on these individual fitness scores. Parental gametes are then produced given recombination and mutation rates, before segregating within mating pairs to generate offspring genotypes. The offspring can disperse to the neighbouring population with a given probability. The loop is then repeated for the next offspring generation.

### Individual based framework

We assume organisms are diploid and composed of males and females. Each individual *i* is characterized by a two-dimensional array that represents a pair of homologous chromosomes. Multiple pairs of arrays may also be constructed to allow the characterisation of any number of chromosomes. Similarly, variation in the number of dimensions of the arrays may be introduced to extend this framework to haploid or polyploid organisms. Each element of the array is an integer defining the copy of a given allele at a given locus. Individuals are also classified into populations. We assume random mating within a population, although this assumption can easily be relaxed (Schindler *et al*. 2015; Ellner, Childs & Rees 2016). Populations *i* and *j* are linked by migration rates (*m*_*ij*_ and *m*_*ji*_) describing movement from population *i* to *j* and vice versa. We assume that individuals that migrate and reproduce successfully pass their genes into the other population hence incorporating gene flow into the model. The genotype-phenotype and phenotype-demography map can differ between populations if required.

The model proceeds in discrete time steps representing generations. It is a forward simulation that includes reproduction and migration at each time step. Density dependence regulates the population growth rate, influencing the probability of successful reproduction (Fig. 1a).

The fitness of individuals is associated with a phenotype (*z*). We only focus on an additive genetic genotype-phenotype map here, but maps including epistasis, pleiotropy and dominance are possible. In the additive case, the sum of values of alleles at each locus gives a breeding value (*b*_*v*_) for each individual at that locus. The sum of breeding values across loci gives a breeding value for the phenotype. Therefore:

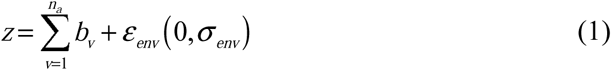

Where *n*_*a*_ is the number of additive loci. In our simulations, the environmental contribution (*ε*_*env*_) is assumed to be stochastic and normally distributed, with mean *0* and standard variation *σ*_*env*_. *ε*_*env*_ may also be dependent on population density or any other environmental driver (Coulson *et al*. 2017).

The fitness function *(ω*) defines the phenotype-fitness map and consequently the type of selection influencing the divergence between populations. Once a population has colonized a novel area, new phenotype-environment interactions appear on the phenotype-demography map, shifting the distribution of phenotypes that are expected to have higher fitness (i.e. phenotypic optima). The difference in phenotypic optima between the populations drives the strength of “divergent selection” (grey area, Fig. 1c). Populations exposed to equal phenotypic optima are considered to be under “concordant selection”. The fitness function we use has the form:

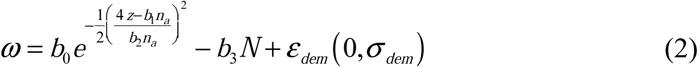

The first part on the right-hand side of equation (2) is based on a Gaussian-distribution determining the relation between the phenotypic trait value (*z*) and fitness (ω). The parameters *b*_*0*_, *b*_*1*_ and *b*_*2*_ define the maximum number of offspring produced, the phenotypic optima, and the variance of the Gaussian curve, respectively. The second part of equation (2) determines the intensity of density-dependence (*b*_*3*_) on the fitness of individuals that are members of a population of size *N*. The final part of the equation introduces a stochastic demographic variant with mean *0* and standard variation *σ*_*dem*_. The last two parts of the equation thus determine the increasing or decreasing variation of fitness due to fluctuations in population size and demographic stochasticity. Any other form of fitness function could be introduced to account for specific relationships between phenotypes (e.g. weight, height, bill size, colour pattern) and the expected number of offspring produced.

The number of breeding events is regulated by the number of females present in the population. Males are randomly selected according to the number of breeding females. The genotypes of both parents participate in the genetic architecture of their offspring by transmitting a haploid copy of genetic material. The offspring differs from the parents by carrying half of the genome of each parent and by specific rules defining the recombination rate (*θ*) between homologous chromosomes. We do not explore the effect of new mutations here, because we are primarily interested in the emergence of genomic islands at relative early stages of evolutionary diversification. However, mutation can easily be incorporated by generating a novel polymorphism at a random locus at a given rate per generation (see example in Appendix S1).

The genetic variation of the new generation is determined by the recombination rate during the segregation of haploid gametes of each parent. Segregation starts with a randomly selected copy of a chromosome (i.e. one of the two dimensions of the individual array defining its genotypes). The recombination rate may either be a fixed value between neighbouring loci or may vary depending on position of the chromosome, for example, through the use of a randomly distributed Poisson process determining crossover points. In the first case, when a recombination map is available, a vector of *n*_*L*_*-1* elements has to be supplied with the recombination rate (*θ*) between each pair of neighbouring loci. The probability of having a crossover (1) or not (0) is uniformly distributed at a rate defined by the value of *θ* between loci (i.e. positions with a probability smaller than *θ* recombine). The uniform distribution allows each position with the same values of *θ* to have an equal chance of crossover across all iterations. There is no recombination between homologous chromosomes when *θ* = 0, both loci are completely linked (e.g. within an inversion or situated close to centromeres), while with a value of *θ* = 0.5, the recombination rate is completely random (i.e. both loci are very distant on the same chromosome or are located on different chromosomes). A value of *θ* < 0.5 means the loci are physically linked. In the second case, a single average recombination rate for the whole chromosome or part of the genotype of interest has to be supplied, and the crossover points are selected by following a random distribution (e.g. exponential). This last method may be preferred when trying to fit a large dataset of genomic information with an unknown recombination rate between neighbouring loci (e.g. Single Nucleotide Polymorphisms). Because we are primarily interested in the effects of various levels of linkage disequilibrium in the formation of islands of genomic divergence, we present results using the first approach, but an example with the second method is also shown in supporting information (Appendix S1).

The offspring represent individuals with the potential to reproduce in the next generation. We assume an equal sex ratio at birth and assign the sexes to offspring by sampling with replacement, with an equal probability of assignment to each sex. A weighted probability could be supplied when unequal sex ratios are considered in the simulations.

The final step is the migration of offspring to neighbouring populations. The probability of migration of each individual is obtained from a uniform distribution, so each individual has the same expected probability of migration. The final number of individuals of population *i* dispersing to population *j* is defined by the migration rate *m*_*ij*_. Individuals of *i* having migration probability smaller than *m*_*ij*_ move to population A value of *m*_*ij*_ = 0 means no migration and thus no gene flow between populations, while a value of 0.5 means random migration (and hence random reproduction) between them.

The final number of individuals in population *i* at time *t+1* can be estimated as the sum of fitness value of all females present in the population at time 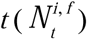 and the number of migrants from population *j* (males and females, 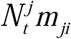):

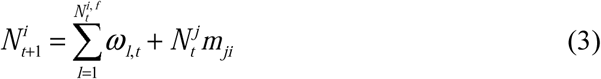

The model is implemented in *R* (R Development Core Team 2017), with some functions written in C++ and integrated to *R* by using the Rcpp package (Eddelbuettel *et al*. 2011). The script is available in the supporting information (appendix S1) and on GitHub (https://github.com/eriqande/gids), and is easily modifiable for further applications. Below we describe a number of simulations with different parameterizations to explore how the signatures of genomic divergence are generated by various processes.

### Initialization

We start by simulating how two populations of diploid individuals with equal intra-genomic variation at the beginning of the simulations diverge. The migration rate between the two populations was varied across different simulations to explore divergence without gene flow (i.e. *m*_*ij*_ = 0) and divergence with gene flow (i.e. *m*_*ij*_ ≠ 0). The demographic and genetic parameter values were chosen to describe two fitness functions that can either have identical or contrasting phenotypic optima, but with a similar number of individuals in each population during the simulation (Fig 1c, Fig 1d, Table 1).

**Table 1.**
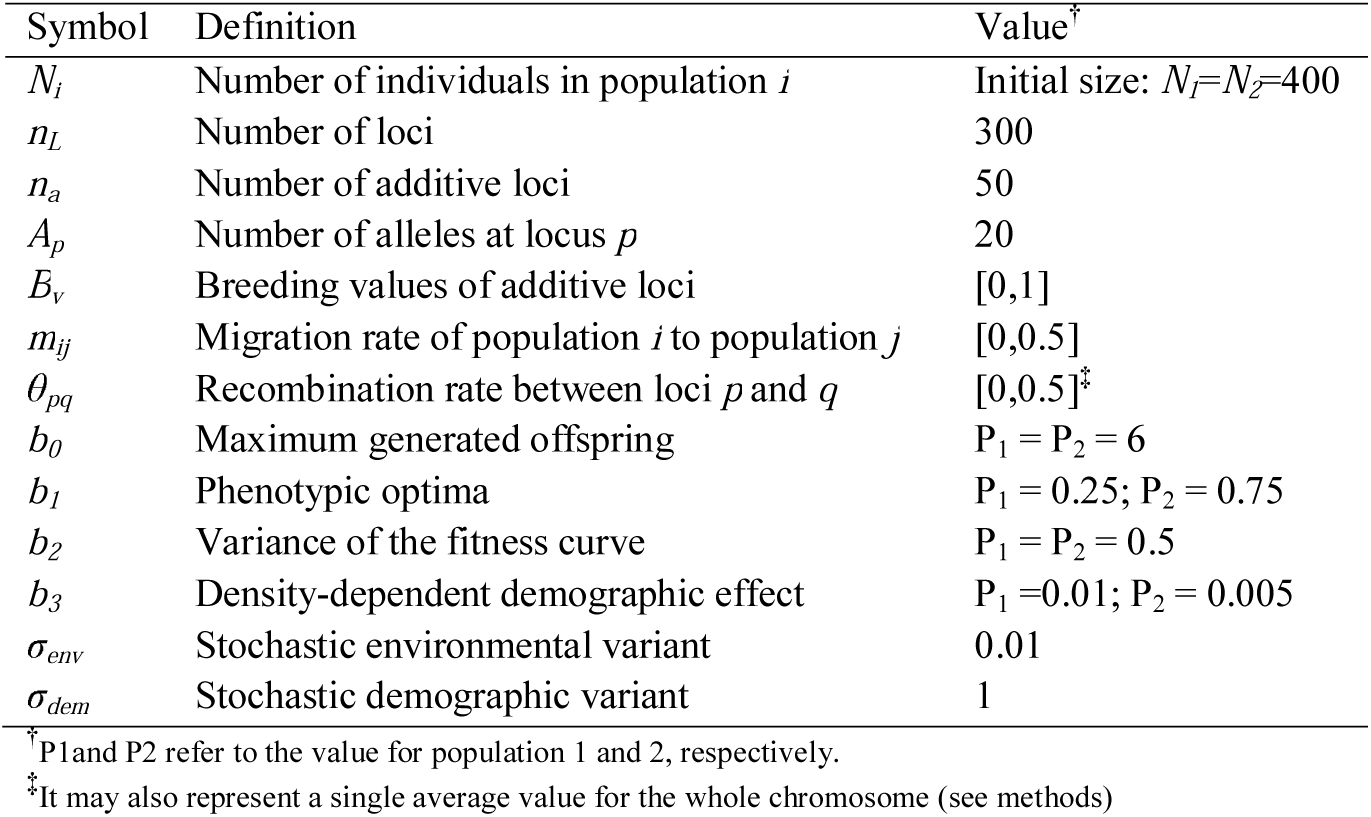
List of parameters of the model with default values

The mean population sizes of the two populations were always around 400 (Fig 1d). This is also the initial number of individuals at the beginning of the simulations. The genomic architecture of individuals was characterized by genotypes across 300 loci (*n*_*L*_ = 300), that were either strongly linked (*θ* = 0.0001) or completely unlinked (*θ* = 0.5). This range of linkage allows us to explore the dynamic of genomic landscapes across more contiguous or distantly related loci. Because previous simulations on the formation of genomic islands of divergence were restricted to bi-allelic loci (e.g. Feder *et al*. 2012; Flaxman, Feder & Nosil 2013), we ran simulations with a higher number of alleles to allow for greater allelic variation (Table 1). The genomic identities of individuals were randomly assigned at the beginning of each simulation by setting the seed of the random number generator in *R*.

Fifty additive loci were chosen to have non-zero variation in allelic contributions to the phenotypic trait value. This fraction of loci is potentially subject to selection. By operating on the phenotype, selection changes the distribution of genotypes at each locus that contributes to the phenotype in the simulation. Loci not influencing phenotypes are neutral and were used to examine the effect of drift and linkage on the appearance of genomic islands. This allowed us to account for both adaptive and neutral evolution simultaneously. The phenotypes were always computed from 50 additive loci (*n*_*a*_ = 50), 10 of which were always linked. These 50 additive loci contributed to the phenotypic trait values of individuals, with the additive value of each allele ranging between 0 and 1. The sum of additive values was then used to compute the phenotype, and then the fitness score, for each individual. However, further studies may expand this procedure to include any required genotype-phenotype map. In summary, we have four classes of genes: i) unlinked genes contributing to the phenotype; ii) linked genes where both loci contribute to the phenotype; iii) unlinked genes that do not contribute to the phenotype; and iv) linked genes that do not contribute to the phenotype. The first two categories of genes are under selection, and the last two are not.

The number of mating pairs depends on the number of breeding females. Female reproductive success was determined first, before male mates were assigned to father each offspring. In this simulation we assumed random mating, although other mating patterns are possible (e.g. Schindler *et al*. 2015). Offspring sex was assigned randomly, with probability 0.5 (Table 1)

### Genomic divergence

We measured pairwise *F*_ST_ at each locus to estimate genetic differentiation between populations. *F*st is a widely used measure of heterogeneity across divergent genomes in studies of genomic islands of divergence (e.g. Ellegren *et al*. 2012; Kusakabe *et al*. 2017). We computed *F*_ST_ at each simulated locus using the *R* package “*pegas*” (Paradis 2010). Genetic differentiation averaged across multiple loci was calculated using the approach of Nei (1973), as implemented in the *R* package “*mmod*” (Winter 2012).

### Simulations

We conducted a number of simulation experiments using a wide suite of parameter values. These were designed to examine how various scenarios of linkage between loci, drift, selection, and time since divergence influence the formation of genomic islands of divergence in both the presence and absence of gene flow. Parameter values are presented in Table 1 and the supplementary information provides more details for the choice of each parameter set (Table S1). The first simulations characterise the effect of the founder population on the resulting genetic divergence (*F*st) at an early stage of independent evolution (100 generations, *m*_*ij*_ = 0). We then explored in more detail, the effect of linkage, gene flow and time since initial divergence. Drift is included in all simulations through the group of genes that are not involved with the phenotype trait value, and through the random selection of gametes at birth. We assigned a name to each group of simulations and will briefly describe their structure.

#### 1. Random sampling of founders and concordant selection

these simulations were designed to examine how random sampling of the founder population influenced genomic divergence. Both populations were exposed to neutral evolution and concordant selection, with identical phenotypic optima (equal to population 1, Table 1). We ran 50 simulations with different random initial founder genotypes. Founder genotypes were determined by sampling a uniform distribution with replacement.

#### 2. Random sampling of founders and divergent selection

We considered the same 50 founder populations as before but added divergent selection. The selective pressures generating evolutionary divergence between populations were generated by their respective fitness functions. The amount of difference between phenotypic optima measures the strength of “divergent selection” (Fig 1c, Table 1).

#### 3. Levels of heterozygosity in the founder population

Our third set of simulations was designed to explore how variable levels of heterozygosity among founder populations influenced the variance of genomic divergence at the end of the simulation. The level of heterozygosity in the founder population was varied by sampling alleles at a locus with variable frequencies of replacement (see Table S1 for more information). The variable frequency of replacement represented the weighted probability of a random sampling with replacement among the 20 polymorphisms available for each locus. This ranged from 1 (an equal probability of allelic sampling and more heterozygous) to 100 (an unequal probability of allelic sampling and more homozygous). As this value becomes higher, it increases the probability for individuals to carry the same allele on both copies of their genes. Because linked loci are hypothesised to be more likely to be involved in the formation of genomic islands and we are analysing this factor separately, we excluded these loci in the final estimation of variability of genetic differentiation.

#### 4. Genomic linkage

Having characterised how initial conditions might influence results, we next examined the effect of linkage on the formation of islands of genomic divergence. We ran 100 simulations with equal founder populations, but changed the recombination rate between linked loci, ranging from nearly complete linkage (θ = 0.0001) to no linked loci (θ = 0.5).

#### 5. Strong selection at a single, unlinked locus

We next explored the effect of strong selection on unlinked genes of large phenotypic effect. Fifty additive loci contributed to phenotypic expression, but one locus contributed 10 times more than the others. This means that rather than having multiple linked loci affecting the trait there is, in particular, one locus of very large effect that is unlinked to the other loci that influence phenotype. We considered the same founder population as in 4 (genomic linkage).

#### 6. Time since divergence with and without gene flow

To explore how time since divergence influenced the formation of genomic islands, we ran simulations with the same 50 founder populations of our previous analysis “random sampling of founders and divergent selection”, but for different lengths of time: 100, 500, 1000 and 2000 generations. We repeated these simulations in the presence (*m* = 0.01) and absence (*m* = 0) of gene flow.

#### 7. Linkage and gene flow

Finally, we explored how linkage and gene flow combined to influence the formation of genomic islands. We simulated various rates of migration and recombination, using the same 50 founder populations as in 4 (genomic linkage). We recorded average *F*st at 10 linked loci affecting the expression of phenotypes (positions 150 to 159) and 10 independent loci not related to phenotype expression (positions 90 to 99). This allowed us to determine the magnitude of differentiation between regions of linked divergent selection and the genomic background of neutral evolution.

## Results

### Random sampling of founders and concordant selection

Our first simulations explored the effect of initial conditions on divergence and the formation of genomic islands under equal selective pressures (i.e. concordant selection). The grey lines in Fig. 2a show the resulting *F*st values for all 50 random initial populations. When considering the average *F*st values by loci across the 50 pairwise comparisons, linked genes that contribute to the phenotype have a slightly higher *F*st than unlinked genes (grey line, Fig. 2b). However, independent of the type of loci (i.e. under selection or neutral), all positions have almost the same probability of becoming an area of higher or lower genomic divergence.

**Fig. 2.**
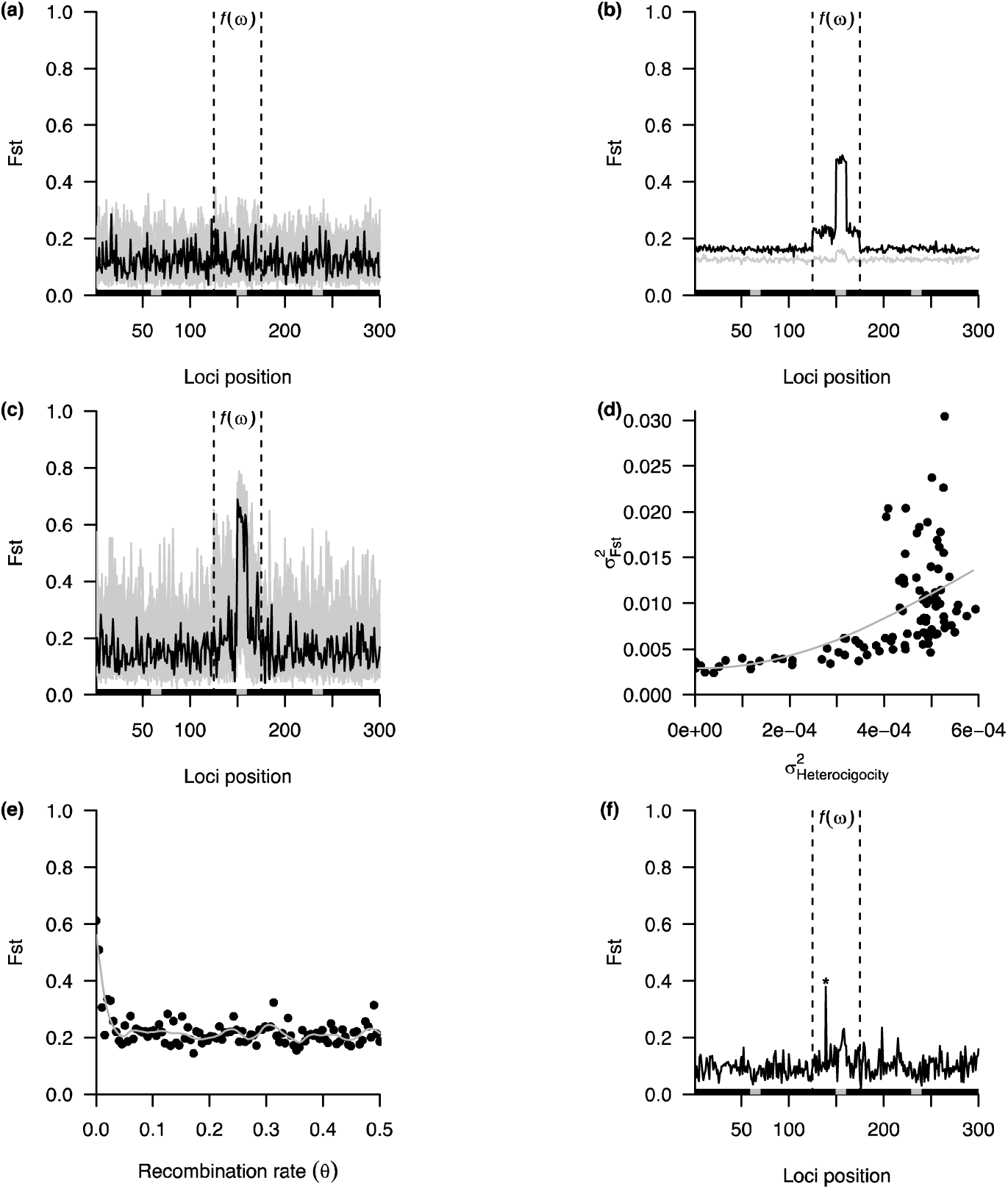
Simulations of genetic differentiation between two simulated populations without gene flow. a) Random sampling of founders and concordant selection. The grey lines represent the resulting genetic differentiation (*F*st) on 50 comparisons with different random sampling of founders. The black line illustrates the resulting values for a single founder population. The grey squares on the horizontal axis represent linked loci (*θ* = 0.0001) and the black rectangles independent ones (*θ* = 0.5). The dotted vertical lines delimit loci participating in the computation of phenotypes. b) Mean divergence by loci on the 50 founder populations. The black and the grey lines represent divergent selection and concordant selection, respectively. c) Random sampling of founders and divergent selection. The grey lines and the black line represent equal founder populations as in figure 2a, but adding divergent selection to the analysis. d) Levels of heterozygosity in the founder population. Influence of the heterozygosity variance at the beginning of the simulations on the variance of *F*st at the end of the simulations. e) Genomic linkage. Effect of the strength of linkage on the formation of a genomic island. *F*st values are averaged over the 10 linked loci influencing the computation of phenotypes and using the same starting conditions as the black line in Fig 2c. f) Strong selection at a single, unlinked locus. A single independent locus with a stronger additive effect on the computation of phenotypes (*). All data are presented after 100 generations of independent evolution.

Different genotypes coding for identical phenotypes influence the dynamics of genetic differentiation with time (Fig. 2a). *F*st values across the whole genome ranged between 0 and about 0.3. Interactions between the genotype-phenotype map and the phenotype-demographic map influence the development of genetic differentiation between populations. The black line in Fig. 2a represents a single founder population with a typical, heterogeneous genomic landscape that has formed over 100 generations. There are areas of higher or lower genomic divergence between the two populations, that appear seemingly randomly across the whole genome. The variance in *F*st we observed within and across loci reveal that the genotype-phenotype map of the founder populations influences the patterns of genomic divergence.

### Random sampling of founders and divergent selection

Genomic islands of divergence are, on average, more likely to be observed for genes that contribute to a phenotypic trait that experiences divergent selection across the two populations (black line, Fig 2b). The range of variance of *F*st values was also higher under divergent selection than under concordant selection (Fig. 2c). The values of *F*st across loci ranged between 0 and about 0.8, and this seemed to affect the average *F*st across non-selected loci (compare grey and black line, Fig. 2b). The same single founder population illustrated in Fig 2a and 2c (black lines) provides an example of where genomic islands form at some linked loci experiencing divergent selection. A single high island of genomic divergence did not emerge at unselected loci. Due to the large variation between simulations, divergent selection did not necessarily generate islands of genomic divergence at loci under selection (grey lines, Fig. 2c).

### Levels of heterozygosity in the founder population

As the variance of heterozygosity in the founder population increases, so too does the variance in *F*st across the genome after 100 generations of independent evolution (Fig. 2d). This variance reflects an increase in *F*st of loci not under selection. *F*st at these loci can be as large as for genes under direct selection. This result reveals that the appearance of a pattern of genomic islands at early stages of differentiation can be caused by the genetic variation at specific loci in the founder populations.

### Genomic linkage

We ran simulations with the same founder population and parameter values used to generate the black line in Fig. 2c that resulted in an island of genomic divergence, except now we varied the recombination rate (*θ*) among linked selected genes. The average *F*st of those linked genes was much higher with nearly complete levels of linkage (*θ* < 0.02), but tended to the average value of neutral genes when the recombination rate was higher, even when they were still physically linked (compare Fig. 2e and Fig. 2b). These results show that strong linkage may facilitate the appearance of genomic islands when those genes are affected by divergent selection, even in the absence of gene flow. Extreme genomic linkage therefore tends to increase the *F*st value of genes under selection. The combined effect of divergent selection and linkage is consequently important for the development of genomic islands of larger sizes.

### Strong selection at a single, unlinked locus

The previous simulations revealed that divergent selection on linked selected loci could sometime result in islands of genomic divergence. We therefore next considered a founder population in which an island of genomic divergence formed (Fig. 2c), yet altered the genotype-phenotype map such that one independent locus (*θ* = 0.5) contributed disproportionately to the phenotypic value. This locus resulted in a level of *F*st of more than twice that observed elsewhere in the genome, including on linked selected genes (Fig. 2f). This result reveals that patterns of genomic divergence are not necessarily determined by strongly linked genes of similar effect, but can also emerge when one gene of large effect is linked to other markers.

### Time since divergence with and without gene flow

We extended our 50 previous simulations of “random sampling of founders and divergent selection” by running them for longer (100 to 2,000 generations). Without gene flow, the trend of higher genomic differentiation in selected loci, particularly in genes that are linked, is more evident at early stages of divergence (100 generations, Fig. 3a). The length of time that independent evolution has to act influences genome-wide divergence, masking signals of genomic islands that arise from single or linked loci. The pattern of heterogeneous genomic differentiation is therefore less evident and tends to disappear as the numbers of generations since divergence without gene flow increases (2,000 generations, Fig. 3a).

**Fig. 3.**
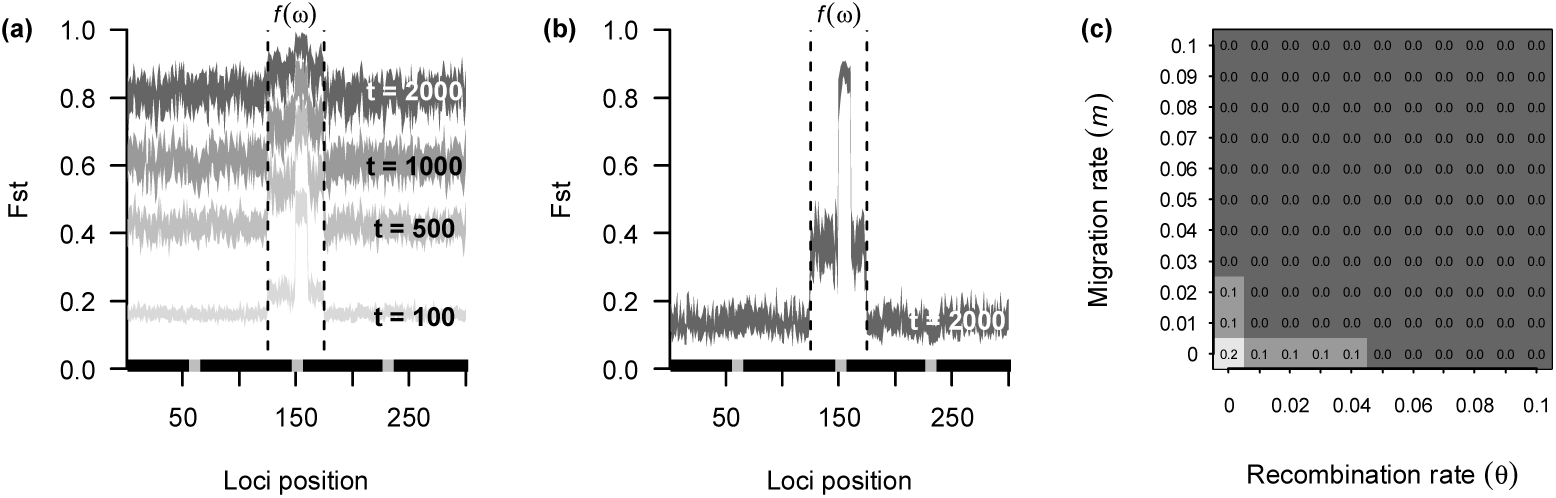
Time since divergence and gene flow. a) Divergence without gene flow. The grey squares on the horizontal axis represent linked loci (*θ* = 0.0001) and the black rectangles unlinked ones (*θ* = 0.5). The dotted vertical lines delimit the loci participating in the computation of phenotypes. The coloured areas represent a confidence interval at 95% of *F*st values, estimated over 50 simulations with randomly assigned genetic identity of individuals at the beginning of the divergence (t = generations). b) Divergence with gene flow. This is similar to the previous figure, but allows for gene flow between populations (*m* = 0.01). c) Linkage and gene flow. Combined effect of migration rate and recombination rate on the magnitude of a genomic island. The numbers inside the squares represent the difference between mean *F*st estimated at the 10 linked loci influencing the computation of phenotypes (i.e. genomic island, positions 150 to 159) and 10 loci not related to fitness and independent (i.e. genomic background, positions 90 to 99). These numbers represent the average difference over the same starting conditions used to estimate the confidence interval of Fig 3a. The data in this last figure are presented after 100 generations of divergent evolution.

In our simulations, populations differentiate with time even in the absence of divergent selection, when both populations have equal phenotypic optima under concordant selection (Fig 2b). This is because there are many ways to generate the same additive phenotypic trait value. The time since initial divergence increases the likelihood of generating these different outcomes (Fig 3a), therefore with enough generations of isolated reproduction, populations can still be highly differentiated even when they are exposed to the same fitness peak.

### Linkage and gene flow

All previous simulations were performed in the absence of gene flow. Gene flow increases the number of generations over which genomic islands of divergence are apparent. The genomic islands of higher *F*st are still present after 2,000 generations, when performing the same simulations as in figure 3a, but allowing a level of migration between populations (*m* = 0.01, Fig. 3b). However, as the level of gene flow increases, the prevalence of islands of genomic divergence decreases (see the zero values in Fig. 3c).

We performed the simulations of divergent selection using the same 50 founder populations (Fig. 2a and 2c), while varying both the migration rate between populations and the recombination rate among linked loci. The numbers inside the grey squares in Fig. 3c indicate the magnitude of difference between independent neutral loci (i.e. genomic background) and linked selected loci (i.e. genomic islands). The largest differences were present under conditions with extreme linkage (*θ* < 0.04) and a low migration rate (m < 0.02), and ranged from 0.1 to 0.2. Those differences are negligible under concordant selection (Fig S2). Overall, these results show that gene flow may influence the persistence of genomic islands but is not the only factor determining their emergence.

## Discussion

### Genomic islands of divergence

The application of our quantitative framework to model the generation of genomic islands of divergence has revealed that while there are several routes that can result in genomic islands, the conditions required to generate islands are relatively narrow, and importantly, there is no single set of circumstances that guarantee their emergence. For instance, formation of large genomic islands requires a combination of divergent selection and strong linkage, regardless of the gene flow scenario. In contrast smaller genomic islands can form via drift in the early stages of divergence in particular. However, in both cases, genomic islands can also fail to form even when these conditions are met, because outcomes are highly dependent on the initial genetic composition of the diverging populations. Our simulations suggest that genomic islands are most obvious during the early stages of divergence, and tend to disappear with the accumulation of genome-wide divergence over time. If present, gene flow can slow this loss up to a point, however including gene flow is not necessary to explain genomic island formation. The importance of evolutionary processes that were modelled (divergent selection, drift, gene flow), along with influencing factors of initial genetic composition, degree of genetic linkage, and time since divergence are summarised in Figure 4. The modelling approach used has provided a nuanced understanding of how genomic islands arise, yet it is not possible to confidently interpret a particular process from genomic data on its own, a long-held goal of genomic data analysis (Turner, Hahn & Nuzhdin 2005; Nosil 2008; Feder *et al*. 2013; Seehausen *et al*. 2014; Nosil *et al*. 2017).

**Fig 4.**
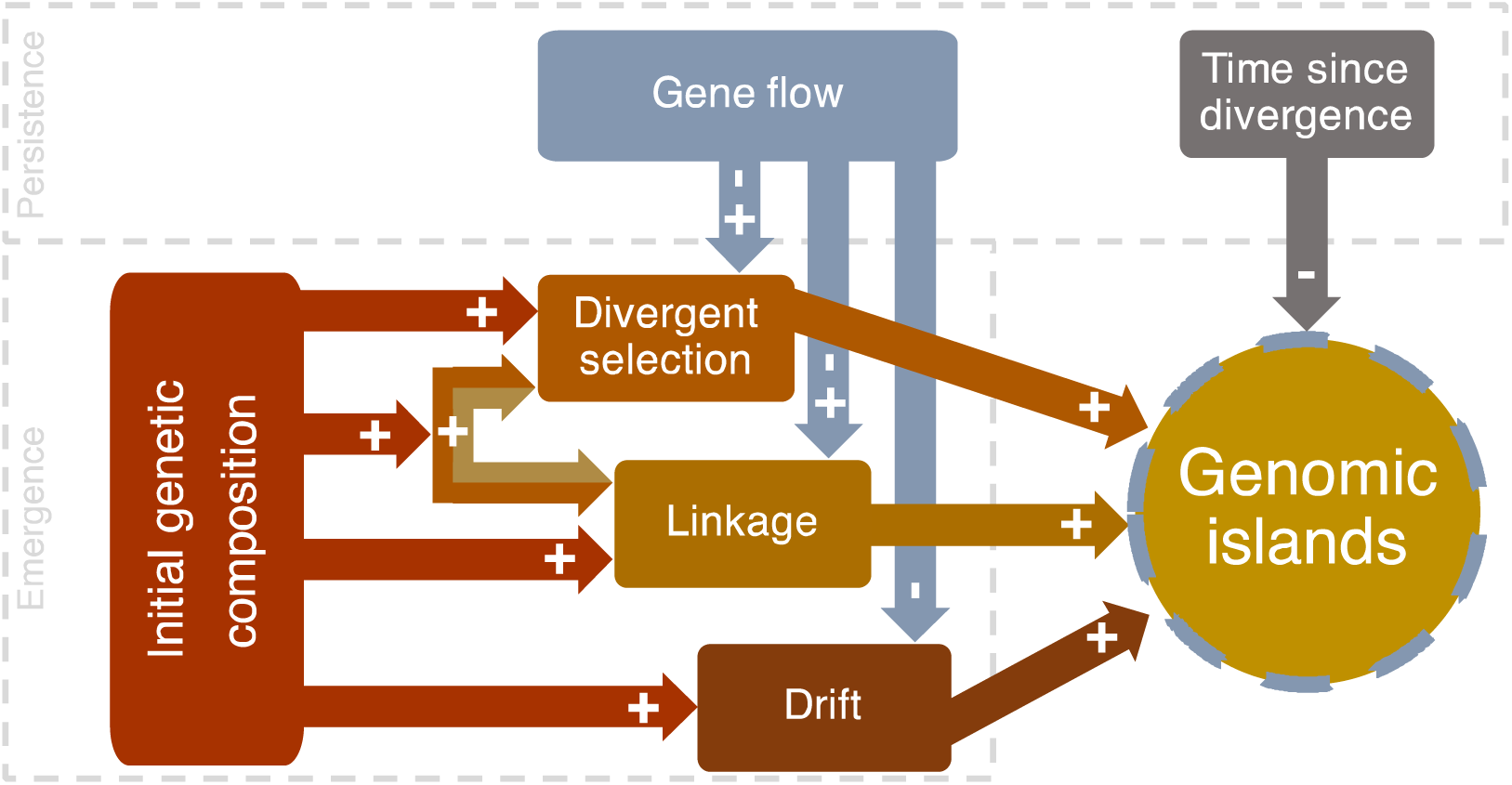
Factors influencing the patterns of genomic islands of divergence. Genomic islands may emerge under the influence of linkage, divergent selection, an interaction between these two factors, or drift depending upon the initial genetic composition of the starting populations (positive effects). Gene flow and time since divergence have an effect on the persistence of islands once formed. Gene flow has an indirect effect by interacting with factors influencing the emergence of this pattern. At early stages of divergence, gene flow can lengthen the time that genomic islands are visible (positive effect), but too high a level of gene flow can erase genomic island patterns (negative effect). The time since divergence has a negative effect on genomic islands, which are more visible under earlier rather than later stages of genomic differentiation.

### Initial genetic composition and drift influence the generation of genomic islands

Our model revealed two ways that random effects can influence the formation of genomic islands of divergence. First, a previously unappreciated but critical factor influencing their generation was the genetic composition of the initial populations. This was evident from comparison of simulations with identical parameter values but different starting populations i.e. different genetic composition. In some simulations, genomic islands were generated and in others they failed to form. Furthermore, the starting values influenced island appearance even in regions of the genome that were not influenced by selection, linkage or gene flow – all of which are thought to be important in genomic island formation as discussed below (Feder *et al*. 2013; Flaxman, Feder & Nosil 2013). Second, drift alone could generate a pattern of numerous islands of small size particularly in the early stages of divergence. Recent studies exploring the distribution of genomic islands have also advanced the idea that islands arise from neutral processes without a major contribution from divergent selection (Campagna *et al*. 2015; Wang *et al*. 2016) and the results of our model identify the scenarios where this is particularly likely to be the case. Some studies document a small number of very prominent islands (e.g. Turner, Hahn & Nuzhdin 2005; Wang *et al*. 2016), however finding multiple islands of low relief is also common (e.g. Ellegren *et al*. 2012; Ruegg *et al*. 2014; Soria-Carrasco *et al*. 2014; Feulner *et al*. 2015). Furthermore, comparisons often involve recently diverged populations (e.g. Nadeau *et al*. 2012; Via 2012; Ruegg *et al*. 2014), with some divergence timescales as short as 100 generations (e.g. Marques *et al*. 2016). Our modelling suggests that these patterns and types of comparisons could be explained without recourse to explanations that invoke selection.

Another scenario that may be particularly prone to stochastic effects is where one of the diverging populations experiences a geographic expansion. Klopfstein, Currat and Excoffier (2006) suggest that the effect of drift is stronger in expanding populations because of “allelic surfing”, where alleles that happen to be at the expansion front may incidentally increase in frequency (see Hofer, Foll & Excoffier 2012; Excoffier, Quilodrán & Currat 2014). This, in turn, impacts genetic composition, and if occurring very early during the divergence process, the combination of early differences in genetic composition and drift could generate highly stochastic patterns of islands of divergence.

We have shown that populations diverge through time even under the equal selective pressures of concordant selection. Indeed, highly polygenic traits may also express divergence based on which alleles of the genes under selection in the founder population end up increasing in frequency. Selection will tend to create shorter coalescence times around those selected loci, meaning a lower effective population size and hence greater drift (Nordborg 1997). However, it should be noted that the specification of the additive genotype-phenotype map we use means there are multiple genotypes that will produce the same phenotypic value. This explains why populations with identical selection regimes can diverge, with some developing islands of genomic divergence, and others not. The nature of our genotype-phenotype map in the model could also underpin the influence of initial genetic composition on our results. Future work will explore whether the same conclusions hold with genotype-phenotype maps that do not assume small additive contributions to the phenotype from genotypes at multiple loci. However, the genotype-phenotype map we use is widely assumed in quantitative genetics, and given that many traits are highly polygenic, is an appropriate initial map to assume in simulations.

### Linkage and divergent selection generate islands of divergence independent of gene flow

Extreme linkage in combination with divergent selection was necessary, though not sufficient, for the development of the most prominent genomic islands, regardless of whether gene flow occurred or not. These findings are consistent with observations of prominent genomic islands between populations presumed to be under strong divergent selection, and not connected by gene flow (Burri *et al*. 2015; Zhang *et al*. 2017). The occurrence of candidate genes, hypothesised to be under natural selection, associated with genomic islands of divergence also supports the role of selection (Sousa & Hey 2013; Kusakabe *et al*. 2017). However, empirical results also provide examples where SNPs under selection are not associated with islands of divergence (e.g. Ruegg *et al*. 2014; Han *et al*. 2017; Riesch *et al*. 2017).

The importance of linkage in the appearance of genomic islands has been highlighted in both theoretical and empirical studies (Feder & Nosil 2010; Renaut *et al*. 2013; Flaxman *et al*. 2014), with extreme linkage, such as that found near centromeres or within genomic inversions, often associated with the most prominent genomic islands of divergence (Feder & Nosil 2009; Ellegren *et al*. 2012; Kawakami *et al*. 2014). Selection acting in these zones of low rates of recombination (i.e. linked selection) reduces the effective population size of these genomic regions to a greater degree than in the rest of the genomes, generating genomic islands (Feder & Nosil 2009; Turner & Hahn 2010).

We did not explore the effect of linkage on deleterious variants (i.e. background selection) in our simulations. However, previous studies have shown that selection on both adaptive and deleterious mutations has a similar effect of reducing within population diversity (Nordborg, Charlesworth & Charlesworth 1996; Slatkin & Wiehe 1998), and influencing the formation of genomic islands (Cruickshank & Hahn 2014).

### The effect of gene flow on generation and persistence of genomic islands

The idea that genomic islands of divergence were generated primarily by antagonistic effects of divergent selection and gene flow was a favoured explanation until recently (Turner, Hahn & Nuzhdin 2005; Nosil 2008; Feder, Egan & Nosil 2012). According to this mechanism, genomic islands form around selected loci involved with the divergence process, and genes physically linked to them, while adjacent neutral or weakly selected regions are homogenised by gene flow (Turner & Hahn 2010; Flaxman, Feder & Nosil 2013; Kawakami *et al*. 2014). Our modelling provides further support that the presence of gene flow is not an essential condition, however, an additional insight is that when gene flow does occur, it can lengthen the time that genomic islands are visible. Verbal models of changing genomic landscapes over time predicted that genomic islands would disappear with the accumulation of genome-wide divergence over time (Wu & Ting 2004; Nosil 2012; Nosil & Feder 2012). Empirical support that this is indeed the case is provided from studies where genomic islands are more frequently documented in recently diverged versus distantly related taxa (e.g. Nadeau *et al*. 2012; Via 2012; Marques *et al*. 2016). Our results reveal that this dynamic can be moderated by gene flow, where a limited amount of gene flow serves to slow down the swamping of genomic islands over time, whereas large amounts of gene flow tend to erase the pattern of islands of divergence altogether.

### Limitations

The main limitation of our simulation approach lies in the amount of genetic and ecological information required to parameterize it for a field system. Empirical information is needed to identify fitness functions, specify the genotype-phenotype map or estimate rates of migration. Model organisms with short generation time that have been extensively studied in the past represent a source of data for potential application (e.g. Mackay 2014). For non-model organisms, applications may adapt the parameter values from sister species for which information is available. This limitation is expected to become less important in the future as the rapid accumulation of freely available ecological and genomic datasets grow (Jones *et al*. 2008; Ellegren 2014). However, in the absence of sufficient information to parametrize a fitness function, this framework is still useful to elucidate neutral evolution, which can be simulated in the framework by replacing the fitness function with a random distribution (e.g. Poisson) in order to generate the next generation of offspring (see example in Appendix S1). While mutation is not explored here, as we were mostly interested in divergence at relatively early stages of evolution, its incorporation would not likely change any of the patterns observed in this study (see example in Appendix S1).

### Conclusions

We have developed a quantitative framework to explore the dynamics of genomic landscapes and identify how various processes can generate patterns of divergence between populations. Our work builds on previous insights (Charlesworth, Nordborg & Charlesworth 1997; Feder & Nosil 2009; Flaxman, Feder & Nosil 2013; Akerman & Bürger 2014; Sedghifar, Brandvain & Ralph 2016). We have been able to demonstrate that the formation of genomic islands of divergence is not a deterministic phenomenon, but that they can arise via a number of routes. We urge extreme caution in inferring a particular ecological or evolutionary process when a particular genomic pattern is observed. Narrowing down the potential cause of a particular signature will likely require ancillary information beyond the genome sequence or modelling exercises that examine the processes that have the potential to generate such a pattern. The methods described here provide a modelling framework which helps to depict such signatures of past evolution, as well as potential routes of future evolution for any divergent taxa.

