## Supporting information for "The multiple population genetic and demographic routes to islands of genomic divergence"

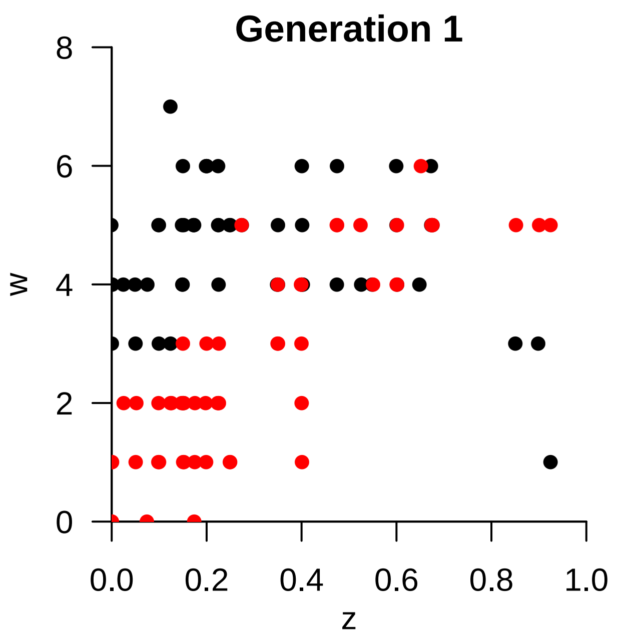


**Figure S1.** Two populations diverging with different phenotypic optima. The fitness ($\omega$) is measured as the amount of offspring and is dependant of an additive phenotype ($z$). Populations are represented with different colour (black or red). Click the picture to start the animation. The animations require Adobe acrobat (reader or professional)

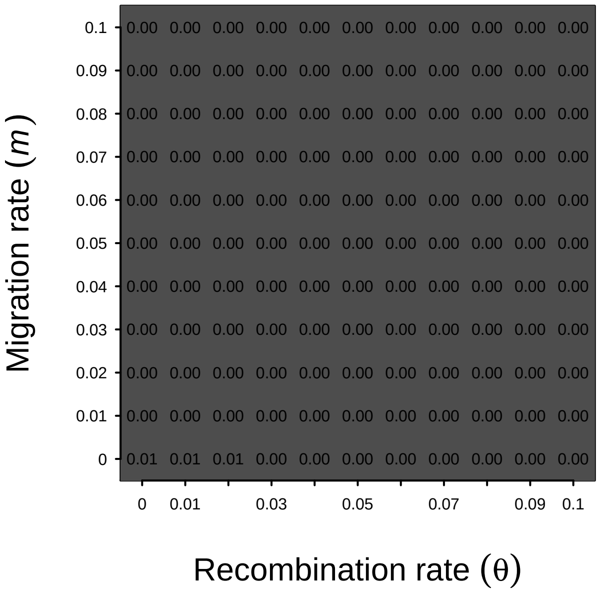


**Figure S2.** Linkage and gene flow under concordant selection. Combined effect of migration rate and recombination rate on the magnitude of a genomic island. The numbers inside the squares represent the difference between mean F*st* estimated at the 10 linked loci influencing the computation of phenotypes (i.e. positions 150 to 159) and 10 loci not related to fitness and independent (i.e. positions 90 to 99). These numbers represent the average difference over the same starting conditions used to estimate the confidence interval of Fig 3a. The data in this last figure are presented after 100 generations.

**Table S1.** Simulated scenarios with parameter values

| **Simulation name** | **N° of simulations** | **$\theta$ linked loci** | $m$ | **Generations ($t$)** | **Prob. alleles**$^*$ | $B_v$ | **Fitness function** | **Figure** |
|---|---|---|---|---|---|---|---|---|
| Concordant selection | 50 | 0.0001 | 0 | 100 | 1 | 0 to 1 | $f(\omega_1) = f(\omega_2)$ | 2a, 2c |
| Divergent selection | 50 | 0.0001 | 0 | 100 | 1 | 0 to 1 | $f(\omega_1) \neq f(\omega_2)$ | 2b, 2c |
| Levels of heterozygosity | 100 | 0.0001 | 0 | 100 | 1 to 100 | 0 to 1 | $f(\omega_1) \neq f(\omega_2)$ | 2d |
| Genomic linkage | 100 | 0.0001 to 0.5 | 0 | 100 | 1 | 0 to 1 | $f(\omega_1) \neq f(\omega_2)$ | 2e |
| Strong selection, unlinked loci | 1 | 0.0001 | 0 | 100 | 1 | 0 to 10$^‡$ | $f(\omega_1) \neq f(\omega_2)$ | 2f |
| Time without gene flow | 1 | 0.0001 | 0 | 2000 | 1 | 0 to 1 | $f(\omega_1) \neq f(\omega_2)$ | 3a |
| Time witht gene flow | 1 | 0.0001 | 0.01 | 2000 | 1 | 0 to 1 | $f(\omega_1) \neq f(\omega_2)$ | 3b |
| Linkage and gene flow | 5000$^†$ | 0 to 0.1 | 0 to 0.1 | 100 | 1 | 0 to 1 | $f(\omega_1) \neq f(\omega_2)$ | 3c |

$^*$ Sampling probability of a candidate allele by locus (1 = equal).

$^†$ The values of $\theta$ and $m$ ranged from 0 to 0.1 by 0.01. We repeated each simulation 50 times.

$^‡$ A single locus has an stronger effect ranging from 0 to 10. Other additive loci range from 0 to 1.
